# Spatiotemporal mapping of immune and stem cell dysregulation after volumetric muscle loss

**DOI:** 10.1101/2022.06.03.494707

**Authors:** Jacqueline A. Larouche, Emily C. Wallace, Bonnie D. Spence, Scott A. Johnson, Mangesh Kulkarni, Eric Buras, Bryan N. Brown, Stephen F. Badylak, Carlos A. Aguilar

## Abstract

Volumetric muscle loss (VML) is an acute trauma that results in persistent inflammation, supplantation of muscle tissue with fibrotic scarring, and decreased muscle function. The cell types, nature of cellular communication and tissue locations that drive the aberrant VML response have remained elusive. Herein, we used spatial transcriptomics integrated with single-cell RNA sequencing on mouse and canine models administered VML. We observed VML engenders a unique spatial pro-fibrotic pattern driven by crosstalk between macrophages and fibro-adipogenic progenitors that was conserved between murine and canine models albeit with varying kinetics. This program was observed to restrict muscle stem cell mediated repair and targeting this circuit in a murine model resulted in increased regeneration and reductions in inflammation and fibrosis. Collectively, these results enhance our understanding of the immune cell-progenitor cell-stem cell crosstalk that drives regenerative dysfunction and provides further insight into possible avenues for fibrotic therapy exploration.

## BACKGROUND

Severe extremity trauma resulting in volumetric muscle loss (VML) incites reductions in muscle function, and quality of life^1,2^. The range and nature of VML injuries as well as alterations in structural and metabolic demands have obfuscated surgical repair schemas and regenerative therapies^3,4^. Currently, the cellular and molecular factors that drive the pathological VML response and prevent healing remain poorly understood. The frank loss of tissue after VML manifests in immune cell infiltration that lasts for days to months^5,6^ and is accompanied by fibrotic scarring^7,8^, which is in contrast with skeletal muscle injury that results in regeneration^9–11^. Previous studies have demonstrated increases in infiltration and accrual of neutrophils^9^, macrophages (mϕs)^6,8,12,13^, and T_helper_ 2^14^ cells in VML injury and the adverse effects of the sustained inflammation from these cells range from exacerbated tissue damage to prevention of muscle stem cell mediated repair. As such, quantitative mapping of immune and stem/progenitor cell dysfunction after VML and how intercellular communication is spatially modified to inhibit regeneration is needed to maximize tissue repair schemas.

Spatial transcriptomics is a novel technology that generates unbiased RNA sequencing (RNA-seq) datasets in a spatially registered manner via capture of transcripts on barcoded beads that are decorated with DNA capture probes and tethered to specific locations on a glass slide^15^. This technique is amenable to integrate with histological imaging, rendering coupled insights into pathological mechanisms of expression changes with spatial context. Recent application of spatial transcriptomics during skin tissue repair^16^ has revealed critical aspects of how injury-responsive cells are recruited to the wound, alter their state and signal with other cell types. Yet, exploration of spatial transcriptomics in the pathological microenvironment of VML injured muscle and elucidating regional variations of immune cell and progenitor functions after VML has not been performed.

Herein, we profiled VML defects using spatial transcriptomics (spGEX) integrated with single cell RNA-Sequencing (scRNA-Seq) datasets to understand the spatial context underlying the development and progression of muscle fibrosis. We observed and validated an abundance of møs localized to the defect zone, mesenchymal derived cells (MDCs) predominantly occupying the transition and defect zones, and myogenic cells primarily localized in the transition zone with negligible infiltration into the defect zone. Cell communication analysis revealed MDCs as key drivers of pro-fibrotic signaling, mediating trophic communication between møs and MuSCs as well as through directly signaling to the møs and MuSCs. Interestingly, time-course spGEX profiling of VML-induced fibrosis in a large animal model revealed co-localization of MDC and myogenic transcripts at an earlier timepoint, which were supplanted by a fibrotic and inflammatory transcriptional landscape by the later timepoint, suggesting conservation of responses across species albeit with different kinetics. Finally, we inhibited pro-fibrotic signaling using a small molecule inhibitor of transforming growth factor beta receptor 2 (TGFBR2) and observed increased infiltration of MuSCs into the defect zone along with reduced inflammatory and fibrotic signaling transcripts across both the defect and transition zones. Together, this work provides a resource for further understanding cell-cell communication networks that contribute to fibrotic degeneration in a spatial context.

## RESULTS

### Spatial transcriptomics of regenerative response to murine volumetric muscle loss injury reveals cellular and molecular pathology

To understand the fibrotic response that develops after VML, we administered 2mm full-thickness punch biopsies to the tibialis anterior (TA) muscles of young adult mice. Consistent with previous results^9^, we observed increased fibrosis by staining with picrosirius red at 7- and 28-days post injury (dpi) (Supp. Figure 1A). To decipher the mechanisms that confer this fibrotic behavior, we extracted cryosections of VML-injured murine tissues from 7-dpi, stained for hematoxylin and eosin (H&E) and annotated the tissue into zones of (1) complete muscle loss (defect zone), (2) remaining in-tact muscle (intact zone) and (3) a transition zone that partitions the lost muscle from the remaining musculature. The defect zone was characterized by an abundance of mononucleated cells consistent with previous observations of inflammation^9^, while the intact zone contained muscle fibers with peripherally located nuclei. The transition zone was characterized by both mononucleated cells, and myofibers with centrally located nuclei indicating active regeneration or degeneration. To glean insights into the complex signaling milieu in the defect and transition zones, we generated replicate spatial transcriptomic maps on VML injured tissues at 7 days’ post injury using the 10x Genomics Visium for spGEX analysis (Figure 1A-B). We generated 247,976,765 sequencing reads, mapped the demultiplexed reads to their corresponding spatial location and observed 2,939 location-specific barcodes with a median of about 13,000 unique molecular identifiers (UMIs) and 3,500 unique genes per spot (Supplemental Table 1). Reads derived from the defect zone displayed more UMIs than the other two zones, which is consistent with the increased number of cells in these locations (Figures 1B-C). We performed gene set enrichment analysis on each zone and observed enrichments in gene sets associated with the immune system, stress, and defense in the defect zone, muscle structure development in the transition zone, and various metabolism gene sets in the intact muscle zone (Figure 1D). Towards this extent, inflammatory (*Ctss, S100a8, S100a9*) and collagen genes were enriched in the defect zone (*Col1a1, Col1a2*), whereas developmental myogenic genes (*Tnnt1, Tnnt2, Myh3, Myh8*) were upregulated in the transition zone, and metabolism genes (*Cox6a2, Cox8b*) and genes associated with mature muscle fibers (*Myh4*) were upregulated in the intact zone (Supp. Fig. 1B). These results demonstrate spatial patterning in gene expression following VML injury is consistent with tissue morphology indicating an area of inflammation, a region of regeneration, and a region that is morphologically intact, but exhibits altered metabolism in response to the defect.

**Figure 1.**
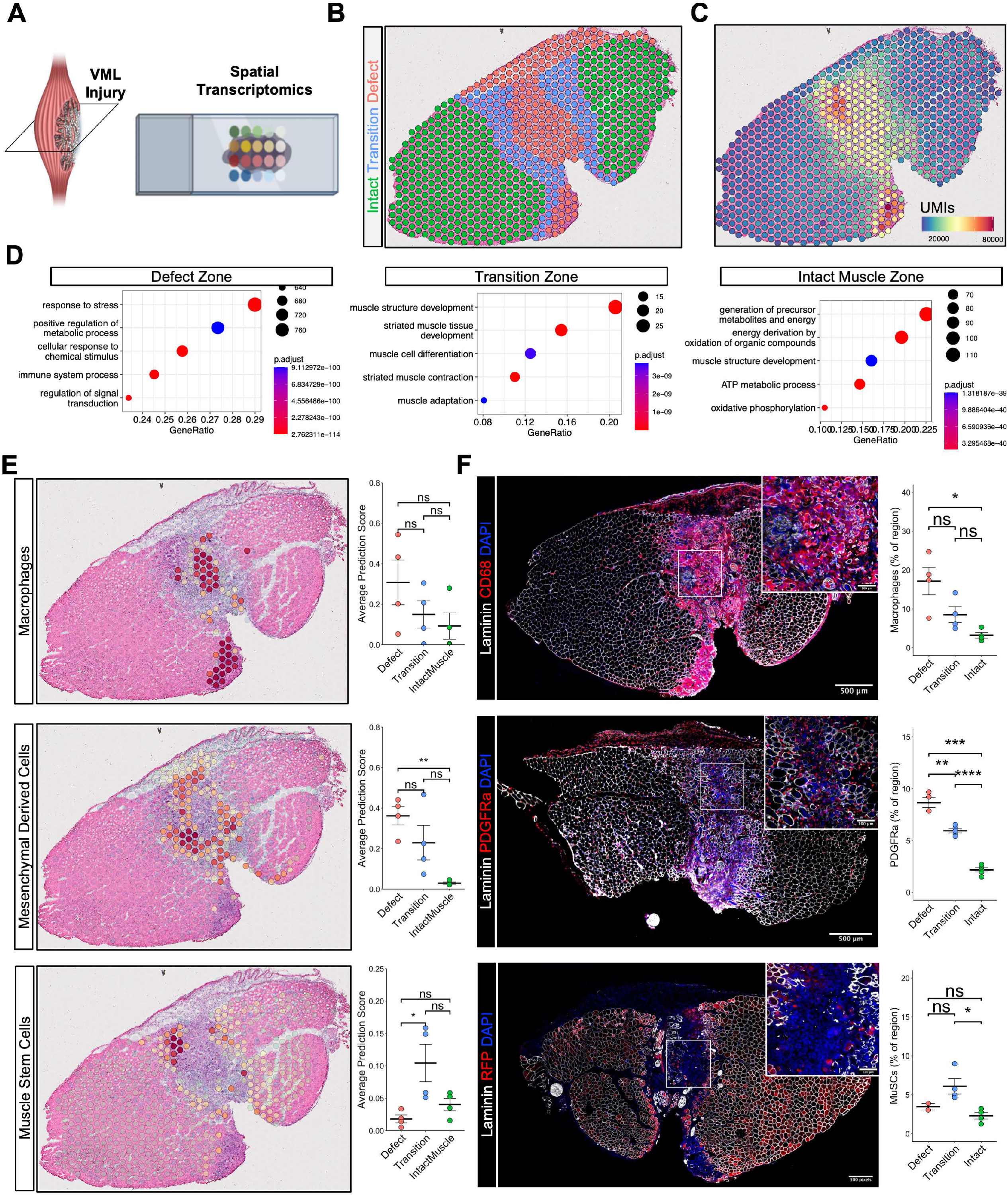
Spatial transcriptomic profiling after volumetric muscle loss reveals pro-fibrotic spatial patterning in the injured site. (A) Experiment schematic whereby spatial transcriptomics was performed on VML-injured tibialis anterior muscles at 7 days post injury. (B) Tissues were annotated into three zones—a defect zone, a zone of intact muscle, and a transition zone between the two. The defect zone is characterized by an abundance of mononucleated inflammatory cells, the transition zone contains centrally nucleated fibers and fibers with small Feret diameters and some mononucleated cell infiltration, and the intact muscle zone contains peripherally nucleated, mature fibers. (C) Distribution of unique molecular identifiers shows higher read counts at the location of the defect. (D) GOTerm analysis on the three zones shows an enrichment in inflammation-associated terms within the defect zone, an enrichment in muscle development and contraction terms in the transitional zone, and an enrichment in metabolism terms within the intact muscle zone. Differentially expressed genes were calculated using Wilcoxon Sum Rank Test with post hoc analysis. log2 fold change > 0.25 and p_adjusted > 0.05 was considered significant. (E) Integration of spatial transcriptomics datasets with matched, cell-type-annotated single cell RNA-sequencing datasets using Seurat label transfer identifies an enrichment of macrophages within the defect and some infiltration into other zones. Pro-fibrotic mesenchymal derived cells are predicted to localize in the defect and transitional zones. Muscle stem cells are absent from the defect zone, but localize in both the transitional zone, and, to a lesser extent, the intact muscle zone. **p<0.01, *p<0.05, and ns denotes p>0.05 by one-way ANOVA with post-hoc analysis. n=4 muscles from 2 male and 2 female mice. (F) Immunohistological stains confirm the spGEX-predictions of cell localization within the different zones. **p<0.01, *p<0.05, and ns denotes p>0.05 by one-way ANOVA with post-hoc analysis. n=3-4 muscles from 2 male and 2 female mice.

A current limitation of spGEX is the low resolution of spatially barcoded spots, which contain reads from up to ten proximal cells^17^. To probe cell localization within our spatial datasets, we used Seurat to integrate our spGEX data with scRNA-seq datasets of VML defects isolated at 7 days post injury that we previously generated^9,18^. This analysis revealed regional localization of cell types consistent with GO Term enrichment analysis and differential gene expression analysis (Figure 1E, n=4 tissues from 4 mice, two-sample, two-sided t-test). For example, the defect zone was predicted to be predominantly occupied by macrophages (møs) and mesenchymal derived cells (MDCs), consistent with the localization of genes such as *Cd68, Ptprc, Aspn, and Col1a1* (Supp. Fig. 2A-B). The transition zone was primarily occupied by muscle stem cells (MuSCs) and their progeny, expressing transcripts associated with myogenesis including *Myog* and *Myh3* (Figure 1E, bottom, and Supp. Fig. 2B). We did not detect expression of MuSC marker genes in the defect zone, which may indicate cellular signaling that inhibits migration or regenerative actions of these cells. To confirm these predictions, we performed immuno-histological staining for CD68^+^ møs and observed nearly identical patterns as our spGEX analysis (Figure 1F, n=4 tissues from 4 mice, one-way ANOVA with BH post-hoc analysis), whereby the defect zone exhibited enrichments in CD68^+^ møs. To validate MDCs, we utilized a lineage-tracing system for PDGFRa^+^ cells (PDGFRa^EGFP^), administered VML defects as above, isolated cross-sections at 7-dpi and performed immunostaining for PDGFRa^+^ møs. Similar to our observations with møs, we detected localized enrichment of eGFP/PDGFRa cells in the defect and transition zones (Figure 1F, n=5 tissues from 5 mice, one-way ANOVA with BH post-hoc analysis). To verify positioning of MuSCs and their progeny after VML, we employed a lineage tracing system for Pax7^+^ cells (Pax7^CreERT2^-Rosa26^TdTomato^) and generated VML defects as above. Cross-sectioning and immunostaining for TdTomato at 7-dpi revealed highly concordant results with our spGEX analysis, whereby MuSCs and their progeny were enriched in the transition zone and were almost entirely absent from the defect region (Figure 1F, n=4 tissues from 4 mice, one-way ANOVA with BH post-hoc analysis). These results suggest that MuSCs are either inhibited from entering the defect zone, or otherwise are unable to remain.

**Figure 2.**
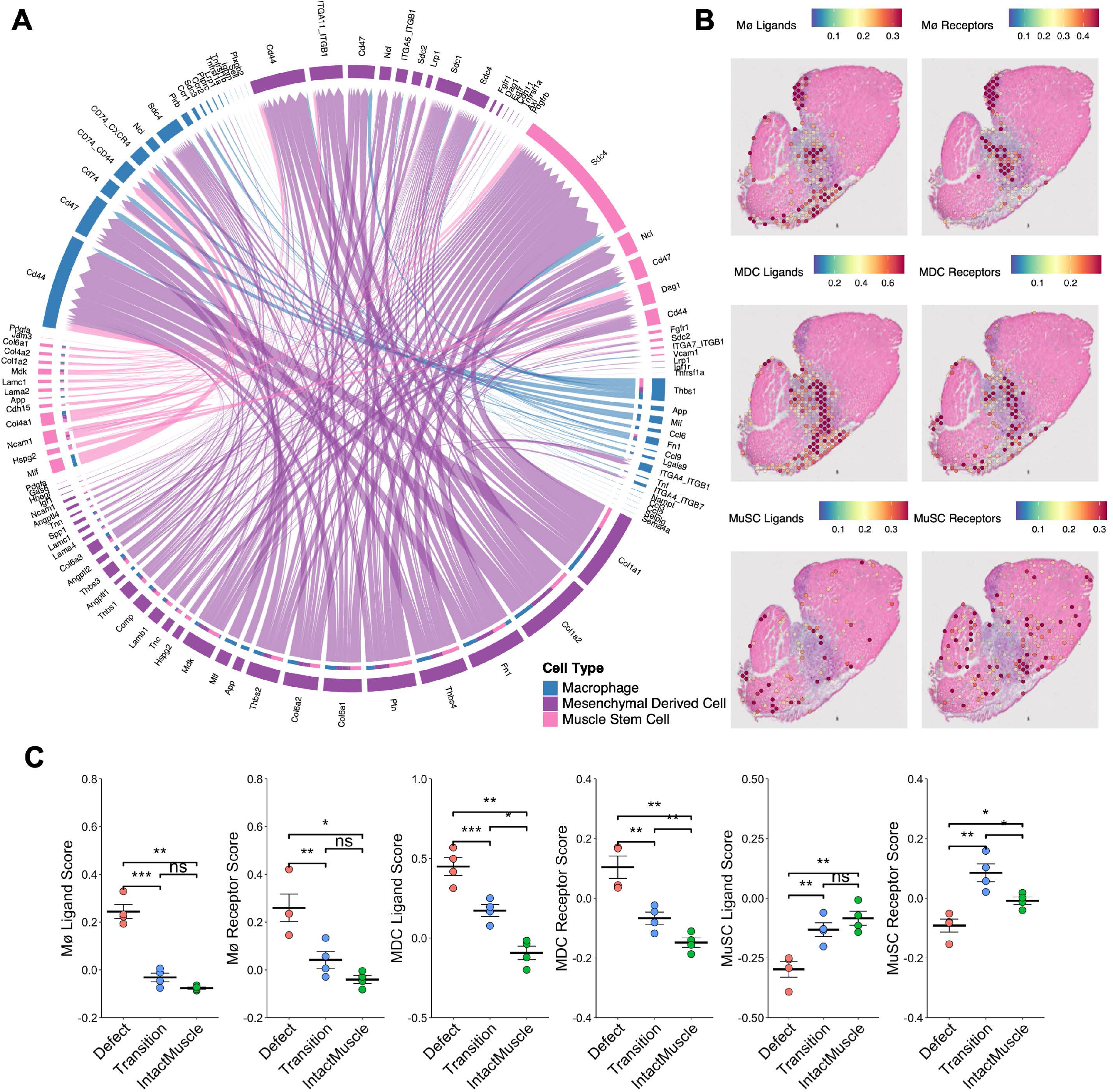
Signaling pathways between macrophages, mesenchymal derived cells, and muscle stem cells post VML are predominantly pro-fibrotic. (A) Chord diagram displaying all significant interactions between macrophages, mesenchymal derived cells, and muscle stem cells determined using CellChat on the single cell RNA sequencing reference dataset. Many of these signaling pathways are associated with fibrosis (collagens, *Comp, Fn1*, laminins, thrombospondins, integrins, *Cd47, Cd44*, and syndecans). Interactions with p < 0.05 based on CellChat’s permutation test were considered significant. (B) Gene module overlays for the ligands and receptors predicted to be involved in significant cell-cell interactions show spatial proximity in the defect and transition zones. (C) Average ligand and receptor module scores across zones demonstrates higher scores for Mø and MDC signaling molecules in the defect zone, increased mesenchymal-derived signaling molecules in the transition zone compared to the intact zone, and higher MuSC module scores in the transition and intact zones. ***p<0.001, **p<0.01, *p<0.05 by two-sided, two-sample t-test. n=4 tissues from 2 male and 2 female mice.

To gain deeper insights into inter-cellular signaling that regulates the regenerative response, we assessed ligand-receptor interactions between predicted cell types in the defect and transition zones. Specifically, we subset the møs, MDCs, and MuSCs from the 7-dpi scRNA-Seq reference dataset and performed CellChat^19^ interaction analysis. We observed substantial crosstalk between the three cell-types, and that the majority of the signaling ligands were expressed by the mesenchymal cells (Figure 2A). Moreover, most of these signaling ligands are pro-fibrotic, including various thrombospondins (*Thbs1, Thbs2*), collagens and extracellular matrix proteins (*Col1a1, Col1a2, Col6a1, Col6a2, Comp, Fn1*). We also observed various inflammatory ligands and receptors, predominantly expressed among the macrophages (*Ccr1, Ccr2, Ccl2, Ccl3, Ccl6, Ccl9, Tnf, Tnfrsf1a, Tnfrsf1b*). To localize these transcripts within the tissues, we calculated module scores for the expression of the CellChat-predicted ligands and receptors and overlaid them onto the H&E images (Figure 2B). This revealed co-localization of the mRNAs for cell-type specific ligands and receptors in the same regions, confirming that the spatial proximity and concentration of these genes in the defect and transition zones supports a pro-fibrotic communication network following VML (Figure 2C, n=4 tissues from 4 mice, one-way ANOVA with BH post-hoc analysis). Together, these results suggest that mø and MPC occupancy of the defect region creates a pro-fibrotic milieu that potentially inhibits MuSC-mediated regeneration.

### Canine VML injury exhibits spatial and molecular homology with murine model

To assess whether our observations from the murine VML model are conserved in a large animal, we administered VML defects to the rectus femoris muscle of female canines by surgical resection of a large volume of tissue (10 cm length × 4 cm width × 2-3 cm depth, Figure 3A-B). The wound was left open and bandaged followed by periodic extraction of punch biopsies (6mm) from different locations of the wound (a more central area labeled as defect and an outer area of the injury labeled as transition). Cross-sectioning, Masson’s staining, and quantitative classification using QuPath^20^ of the extracted biopsies revealed five different types of tissue (debris, healthy in-tact muscle, necrotic muscle, granulation tissue and connective tissue, Figure 3C). At the superficial edge of the wound, we observed a thick scab and granulation tissue at both time points and tissue locations, respectively. At seven days post VML injury, both healthy in-tact muscle and connective tissue were co-located in the deep edge of the wound with necrotic muscle interspersed between the superficial and deep edge. At 14 days post VML injury, development of a fibrotic scar, as observed in our murine model, is detected with a minimal amount of healthy in-tact muscle in the transition location. These results demonstrate a conserved response to VML whereby muscle regeneration is initiated and supplanted by fibrosis.

**Figure 3.**
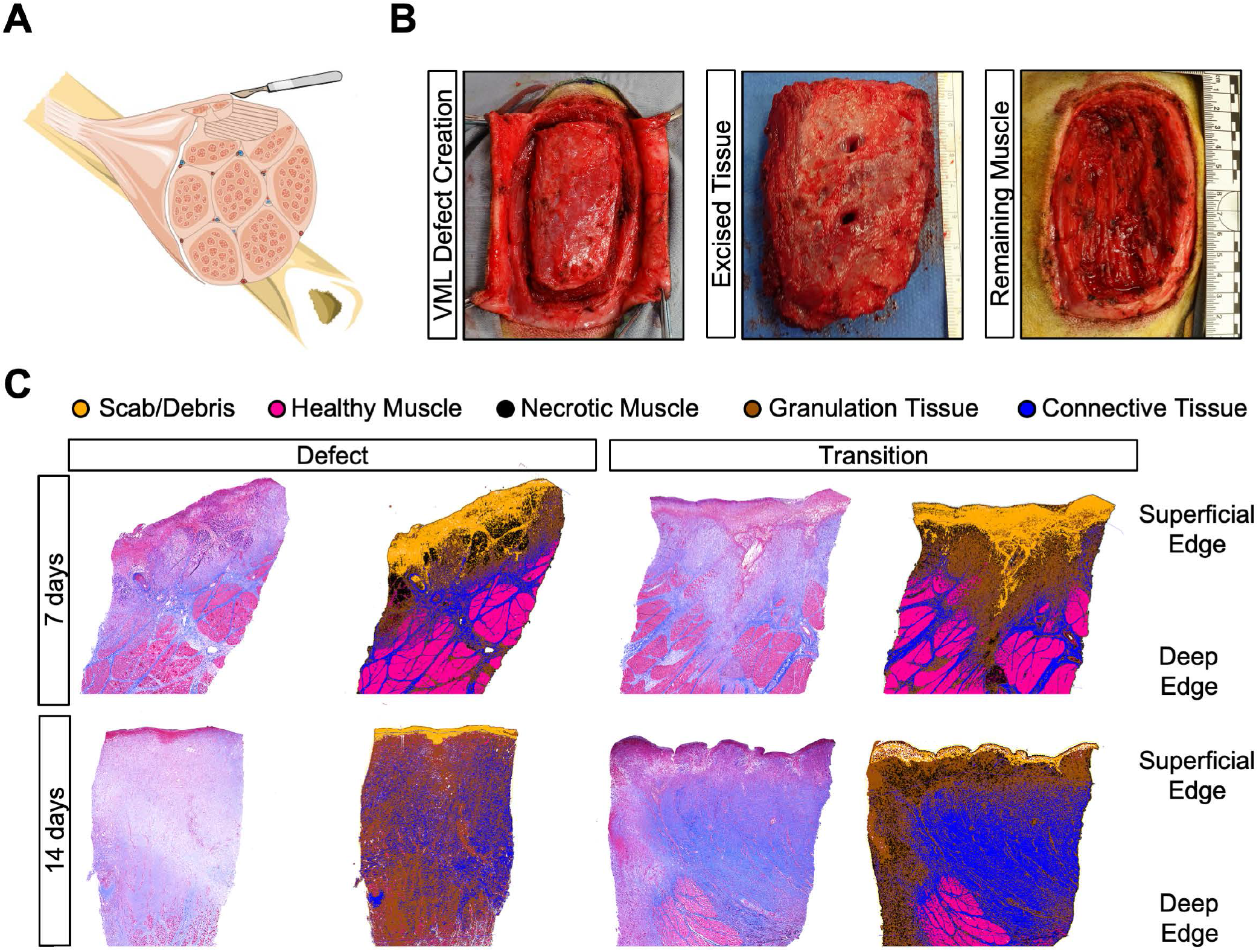
Administration and characterization of VML defect in a canine model. (A) Schematic of volumetric muscle loss injury to canine biceps femoris muscle. (B) Images of VML defect creation whereby the skin is opened and muscle is exposed (left) followed by surgical removal of a 10 cm × 4 cm volume of muscle (middle) and the remaining wound is left open (right). Scales are shown in middle and right images. (C) Masson’s trichrome-stained sections of the punch biopsies. The superficial edge of the wound is located at the top of the image. Images were manually segmented to identify scab/debris (orange) at the wound surface, and automated image analysis using QuPath was performed to automatically identify healthy muscle (pink), necrotic muscle (black), granulation tissue (brown), and connective tissue (blue) within the remaining image. The top 6 mm of the sample, representing the active wound site at 7 and 14 days is shown.

To determine if the molecular observations we made in the murine model were conserved in the canine, we performed spatial transcriptomics on tissue sections collected at 7- and 14-days post VML injury in a canine. Biopsies were collected from both a more central area and an outer area of the injury (Figure 4A). All biopsies, including from both timepoints, were collected from the same animal. At 7-dpi, some remaining myofibers were observed within the defect, whereas by 14-dpi there was a dense region of mononucleated cells at the surface of the defect and substantial extracellular matrix with some cellularity below, indicative of fibrosis (Fig. 4B). We generated nearly 455 million sequencing reads, and each demultiplexed read was spatially mapped yielding 8,636 location-specific barcodes with a median UMIs per spot ranging between 4-8.5 thousand across the tissues, and a median unique genes per spot ranging from 1.5-2.3 thousand across the tissues (Supp. Table 2, Supp. Fig. 3). SpGEX data normalization, scaling, dimensional reduction, and clustering revealed transcriptional similarity among samples collected from the same timepoint, though spatial heterogeneity was also evident as the spots from different tissues clustered separately (Fig. 4C). We performed differential gene expression analysis across the samples and identified transcripts indicative of an initial myogenic response including upregulation of troponins (*TNNT1, TNNT3, TNNC1*) within the defect region at 7dpi. At the outer edge of the wound at 7-dpi, upregulated transcripts were largely associated with inflammation (*CTSS, PLAU, CCL2, CCL3, CCL4, SPP1*). In contrast, at 14-dpi, both tissue samples were upregulated for transcripts associated with ECM deposition (*COL11A1, FN1, ACTG2*) and ECM remodeling (*MMP1, MMP12*), as well as inflammation (*S100A8, IL1R2, PTGS2, S100A12, CXCL8*) (Figure 4D).

**Figure 4.**
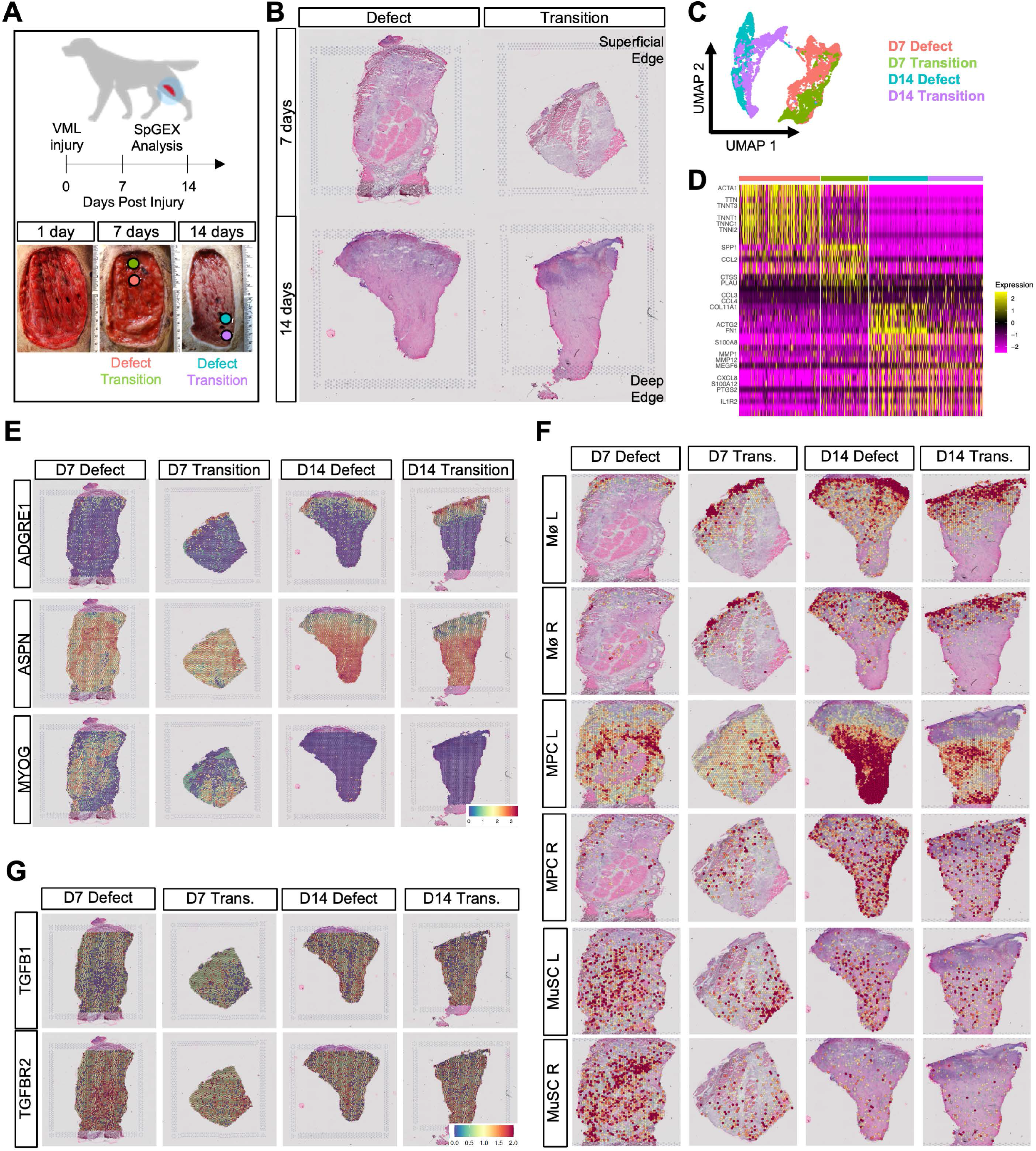
VML in canine results in a regenerative response that is supplanted by a profibrotic network. (A) Schematic of experiment design whereby tissues were collected from the center (defect zone) and edge (transition zone) of the defect at 7- and 14-days post VML injury to canine rectus femoris muscle. 10x Visium spatial gene expression analysis was performed on sections of each biopsy. (B) H&E-stained sections of the punch biopsies. The superficial edge of the wound is located at the top of the image. (C) UMAP dimensional reduction of spots colored by sample. (D) Heatmap of top differentially expressed genes according to sample. (E) Overlays of marker genes for macrophages (*ADGRE1*), mesenchymal cells (*ASPN*), and myoblasts (*MYOG*). Myoblasts gene expression is observed in the defect at 7-dpi, but is completely absent at 14-dpi, whereas mø and MDC associated gene expression increases. (F) Overlays of CellChat-predicted active signaling modules shows high expression at 14-dpi of MDC and mø ligand and receptor genes and reduced expression of MuSC signaling genes. (G) Overlays of *TGFB1* and *TGFBR2* show elevated expression throughout all biopsies at both 7 and 14-dpi.

Next, we probed whether cellular dynamics and spatial patterning observed in the murine model were conserved in the canine VML defects. Overlays of marker genes for møs (*ADGRE1*), MDCs (*ASPN*), and myogenic cells (*MYOG*) suggest myogenic cells and MDCs are present within the defects at 7-dpi, with some møs localized principally on the superficial edge (Figure 4E). By 14-dpi, myogenic gene expression is undetectably low, while expression of both *ASPN* and *ADGRE1* is increased. Moreover, the møs, based on *ADGRE1* expression, appear to infiltrate deeper into the defect (Figure 4E). To assess whether similar signaling pathways were involved in the canine response as in the murine response, we generated gene module scores with the canine gene analogs to the significant LR interactions predicted by CellChat from the murine data (Figure 4F). The pro-fibrotic signaling molecules were highly expressed at 14-dpi, and spatially aligned with marker gene expression for MDCs and møs. We also observed some overlap of mø and MDC signaling genes, suggesting that these cell types are communicating in the canine model as well to contribute to fibrotic development. The expression of myogenic signaling genes was higher at 7-dpi and more localized to where MDC signaling transcripts were expressed than where mø genes were. Since several of these LRs are associated with TGFβ-signaling (*CD47, THBS1*, and others), and TGFβ-signaling is a known driver of muscle fibrosis via FAPs^21^ and MuSCs^22–24^ in rodent models, we sought to understand localization of TGFβ-signaling in the canine. We observed substantial expression of both *TGFB1* and *TGFBR2* throughout the defects at all timepoints (Figure 4G). While *TGFB1* expression is similar at both timepoints, *TGFBR2* is upregulated at 7-dpi, which suggests may participate in the myogenic to fibrotic switch. Together, these results suggest that both the cellular and molecular patterns that occur in the murine VML model are conserved in a large animal model, though the timeline of the response is delayed.

### Interruptions to TGFβ modulates spatial crosstalk that contributes to fibrosis

Based on the observed activation of fibrotic signaling between møs, MDCs, and MuSCs post VML, high expression of *TGFB1* and *TGFBR2* in the canine, and prior observations of improved functional recovery, enhanced myogenesis, and reduced fibrosis post TGFBR2 inhibition in mice^9,23^, we sought to understand the spatial and signaling implications of pharmacologically inhibiting TGFBR2. A cohort of mice received bilateral VML defects to the TA followed by treatment with ITD1 in one limb and vehicle in the contralateral limb (Fig. 5A). At 7-dpi, we stained sections with H&E to observe and annotate tissue morphology into the three zones and generated matched spGEX datasets (Supp. Fig. 4A-B). Consistent with improvements in function and reduced fibrosis, gene set enrichment analysis comparing the defect zone in treated and untreated TAs showed upregulation of terms associated with myogenesis and muscle repair (Supp. Fig. 4C). Integration with scRNA-Seq datasets to localize cell populations across the tissues predicted reductions in macrophages throughout the tissue, reductions in MDCs in the transition zone, and the presence of myogenic cells in the defect zone (Fig. 4B, n=4 tissues from 4 mice, two-sample, two-sided t-test). Immunofluorescent staining of serial tissue sections confirmed the spGEX predictions of reduced PDGFRa+ MPCs and increased myogenic cells following ITD1 treatment, though CD68+ macrophages were increased in the defect zone (Fig. 4C, Supp. Fig. 4D-F, n=3-4 tissues from 3-4 mice, two-sample, two-sided t-test). This result may be driven by transcriptional differences in the macrophage populations that result from ITD1 treatment. Together, this suggests that inhibiting TGFβ signaling via the TGFBR2 receptor reduces the biochemical signaling within the defect zone that is preventing a MuSC-mediated regenerative response.

**Figure 5.**
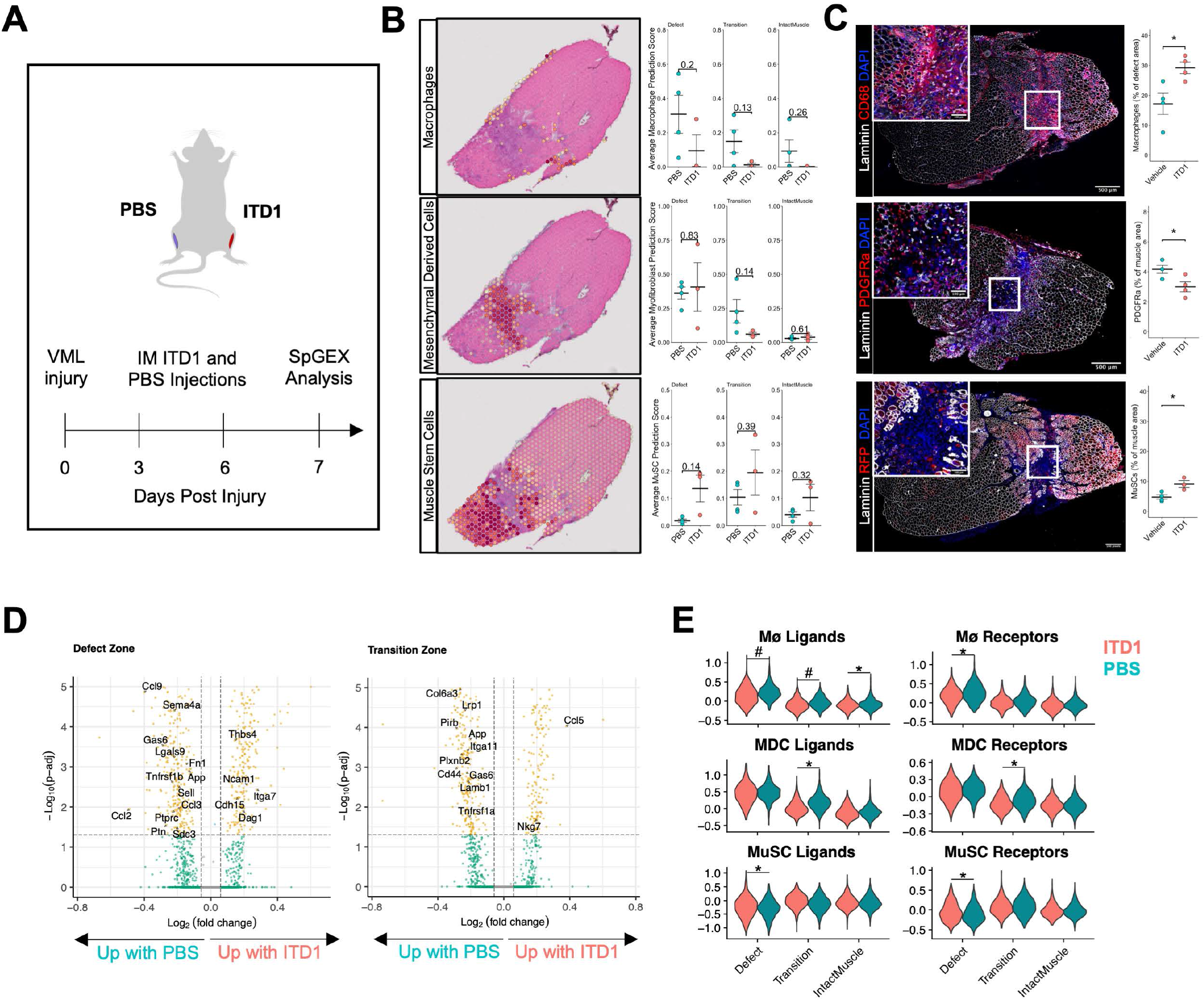
TGFb signaling inhibition reduces inflammation and enhances myogenesis within the VML defect. (A) Experiment schematic whereby a cohort of mice received bilateral VML defects to the tibialis anterior muscles followed by intramuscular injection of saline (PBS) or TGFBR2-inhibitor ITD1. Spatial transcriptomics analysis was performed at 7 days post injury. (B) Integration of spatial transcriptomics datasets with matched, celltype-annotated single cell RNA-sequencing datasets using Seurat label transfer identifies a trend towards reduced macrophages with the fibrotic transcriptional profile in all zones, fewer fibrotic mesenchymal derived cells in the transition zone, and increased numbers of muscle stem cells in the defect zone. Plots are annotated with p-values. n=3-4 tissues from 6 mice. Statistics performed using two-sided, two-sample t-test.(C) Immunohistological stains of macrophages (CD68), PDGFRa+ mesenchymal cells, and RFP+ MuSCs and their progeny using a lineage tracing model show increased macrophages as a percentage of the defect region, reduced PDGFRa+ mesenchymal cells, and increased MuSCs and their progeny following ITD1 treatment. *p<0.05. n=3-4 tissues from 6 mice. Statistics performed using two-sided, two-sample t-test. (D) Volcano plots showing differential gene expression as a result of ITD1 treatment in the defect (left) and transition zones (right). Many of the genes identified by CellChat as activated signaling genes were downregulated with ITD1 treatment. Differential expression was calculated using MAST, and genes with a p-adjusted value less than 0.05 were considered significant (yellow). (E) Violin plots of gene module scores for the macrophage, mesenchymal derived cell, and MuSC ligands and receptors (L&Rs) predicted to be involved in active signaling pathways post VML shows reduced expression of Mø L&Rs in the transition zone, reduced expression of MDC L&Rs in the transition zone, and increased expression of MuSC L&Rs in the defect zone. Though not statistically significant, Macrophage L&R expression was also reduced in the defect zone post treatment with ITD1. *p<0.05, #p = 0.06 by two-sample, two-sided t-test comparing average module scores for each tissue in each zone across treatments. n = 3-4 tissues per group from 3-4 mice.

To further understand the implications of ITD1 treatment on regeneration post-VML, we performed a MAST^25^ differential gene expression analysis on the defect and transition zones (Figure 4D). In line with increased myogenic cells populating the defect, we observed increased *Cdh15* and *Ncam1* expression along with reduced inflammatory signaling genes (*Ptprc, Ccl2, Ccl3, Ccl9, Gas6, Tnfrsf1b, Lgasl9, Sema4a*). Within the transition zone, ITD-1 treatment reduced expression of several of the pro-fibrotic ligands and receptors predicted by CellChat (*Cd44, Tnfrsf1a, Gas6*). Interestingly, among the upregulated genes were *Nkg7* and *Ccl5*, which are associated with NK cells^26^ and cytolytic NK cell signaling post VML^9^. Overall, ITD1 treatment reduced many of the pro-fibrotic signaling ligands and receptors predicted to be activated post VML by CellChat (Figure 4E), including reduced mø ligands in all zones, reduced mø receptors in the intact zone, and reduced MDC ligands and receptors in the transition zone. In line with improved regeneration and an enhanced myogenic response, MuSC ligands and receptors were increased in the defect zone post treatment (n=4 tissues from 4 mice, two-sample, two-sided t-test). Collectively, our results show that blocking TGFβ1 signaling creates a biochemical environment in the defect zone more amenable to the MuSC regenerative response, acting on both the immune and mesenchymal accumulation and localization, and favorably altering signaling networks between møs, MPCs, and MuSCs.

## DISCUSSION

VML has consistently demonstrated a failure of regeneration^27^, fibrotic scarring^5,8^ and reductions in muscle function^2^, but the cellular and molecular mechanisms that confer this behavior and their spatial context remain poorly understood. As a result, many therapeutics for VML have displayed limited improvements in muscle regeneration and functional output^3,4^. Our results address this need and demonstrate a conserved a signaling circuit between møs and MPCs in murine and canine models of VML that contributes to fibrotic development and impinges on MuSC-mediated regeneration. Targeting the cellular crosstalk between møs and MPCs through TGFβ inhibition improved muscle regeneration and infiltration of MuSCs in the defect as well as dampened fibrosis.

The microenvironment is known to impact cellular behavior and understanding cellular neighborhoods and which cell types co-localize in morphological regions that become dysregulated is critical to understanding signaling mechanisms and informing therapeutic development^28^. Herein, we identified a spatial patterning whereby møs and MDCs heavily populate the region of muscle loss post-VML, while myogenic cells inhabit the boundary region between the defect and remaining intact muscle. The absence of MuSCs in the defect region at 7-dpi in the mouse and 14-dpi in the canine may be a result of an inability to migrate into or reside within the defect. We speculate this behavior may be mediated by an inadequate biophysical microenvironment^29^ (increased tension of the matrix or lack of binding sites) and/or an inhibitive biochemical microenvironment^30^ driven by mø-FAP signaling^31^. In support of this, we observed a highly similar program as chronic muscle fibrosis, whereby inflammatory møs secrete latent TGFβ1 that is activated biochemically by FAP-secreted factors and mechanically through strain on a stiffened matrix. The activated TGFβ1 in turn induces FAP differentiation towards myofibroblasts^32^, and reduces TNFα-induced FAP apoptosis^33^. It is likely that the mø-FAP-directed communication is part of a feed-forward circuit whereby the FAPs also recruit møs and monocytes and direct their phenotype towards fibrosis^34^. Co-localization of these two cell types within the defect, the predominantly pro-fibrotic ligand-receptor pairs, and the reduction of FAPs and MDCs observed post TGFβ-inhibition suggests similar mechanisms occur post-VML.

FAPs are also important mediators of MuSC differentiation and self-renewal^35^ following injury, in part due to their secreted factors. In the canine, which contains larger distances over which host tissue must respond to injury, we observed similar cellular dynamics of co-localized møs and MDCs, but varied kinetics whereby higher *PDGFRA* and myogenic gene expression is observed at 7-dpi, followed by the complete absence of myogenic gene expression at 14-dpi and increased expression of myofibroblast genes. Increased expression of *COL1A1* and *THY1* at 14-dpi in the canine also suggests a phenotypic transition of FAPs towards fibrogenesis^36^, which was at least partially ameliorated by ITD-1 treatment. It remains unclear how positioning between møs and MDCs is established in the VML wound and whether contraction of the fibrillary collagen matrix from MDCs signals to møs to promote chemotaxis. Conversely, increased secretion of latent TGFβ1 by møs could drive increases in integrin binding in MDCs and induce matrix contraction. Further research is warranted to deconstruct the recruitment and ordering of cells that drive fibrotic scarring and inhibition of MuSC migration in the VML defect.

Muscle stem cell-mediated regeneration is coordinated by soluble cues from resident and infiltrating immune and progenitor cells, as well as physical structure provided by the extracellular matrix^37,38^. The absence of MuSCs in the defect region at 7-dpi in the mouse and 14-dpi in the canine may be a result of an inability to migrate into or reside within the defect. Our observations that MuSCs have infiltrated into the defect at 7-dpi in the canine but are then replaced by dense ECM and abundant MPC marker gene expression at 14-dpi combined with our observations of more sustained MuSC infiltration into the defect region following TGFβ signaling inhibition suggests that manipulating inter-cellular signaling and the biochemical milieu resulting from VML could enhance regeneration. In line with these observations, we observed increases in expression of NK cell-related transcripts with ITD1 treatment. Prior work from our group has shown that NK cell infiltration into VML injuries reduces neutrophil abundance, and that the neutrophil secretome impairs myogenesis in vitro^9^. Thus, enhanced NK cell activity and restrained inflammation from neutrophils as a result of TGFBR2 inhibition could be one mechanism through which this intervention contributes to a more favorable biochemical environment for muscle regeneration.

VML continues to remain a significant clinical need and our results enhance understanding of the pathological drivers of this trauma. These datasets may yield enhancements in existing regenerative therapies and useful aids for quantifying therapeutic effects for VML.

## Acknowledgments

The authors thank the University of Michigan DNA Sequencing Core for assistance with spatial sequencing library preparation. The authors also thank Josh Welch, Benjamin Levi, and Robert Tower for advice regarding bioinformatics analysis, along with other members of the Aguilar laboratory.

## Author contributions

J.A.L., E.C.W., B.D.S., S.A.J., and M.K. performed the experiments, J.A.L., analyzed the data. J.A.L., and C.A.A. designed the experiments. E.B., B.N.B., and S.F.B. contributed reagents. C.A.A. supervised the work. J.A.L. and C.A.A. wrote the manuscript with additions from other authors.

## Figure Captions

**Supplemental Figure 1.**
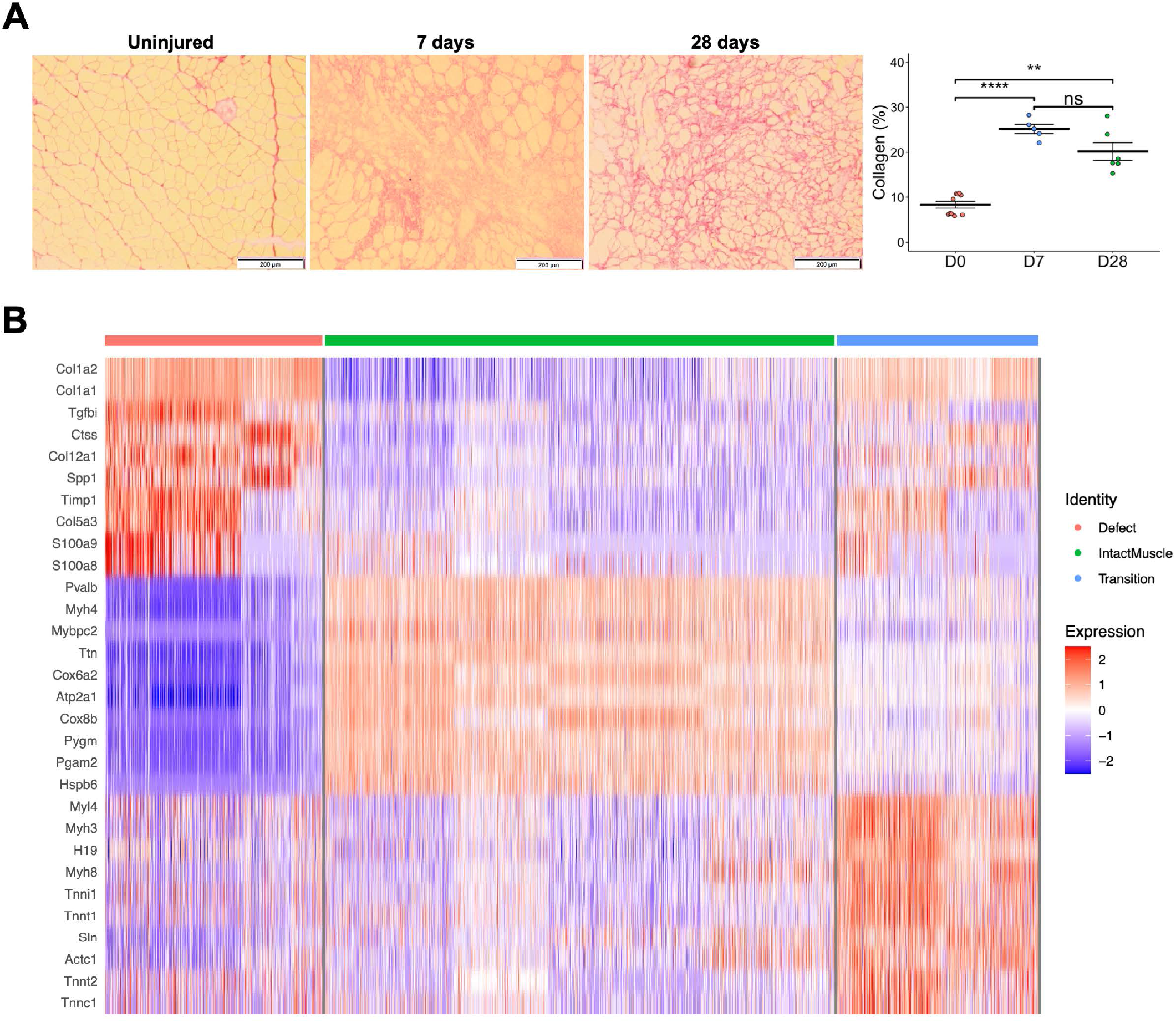
Characterization of volumetric muscle loss in tibialis anterior muscle. (A) Representative images of tissues stained with picro-sirius red as well as quantification of collagen at each time point. n = 6-10 muscles. Full section 10X stitches were analyzed for each tissue. **p<0.01,****p<0.0001 by two-sided one-way ANOVA and Tukey’s post-hoc analysis. (B) Heatmap of gene signatures for each zone.

**Supplemental Figure 2.**
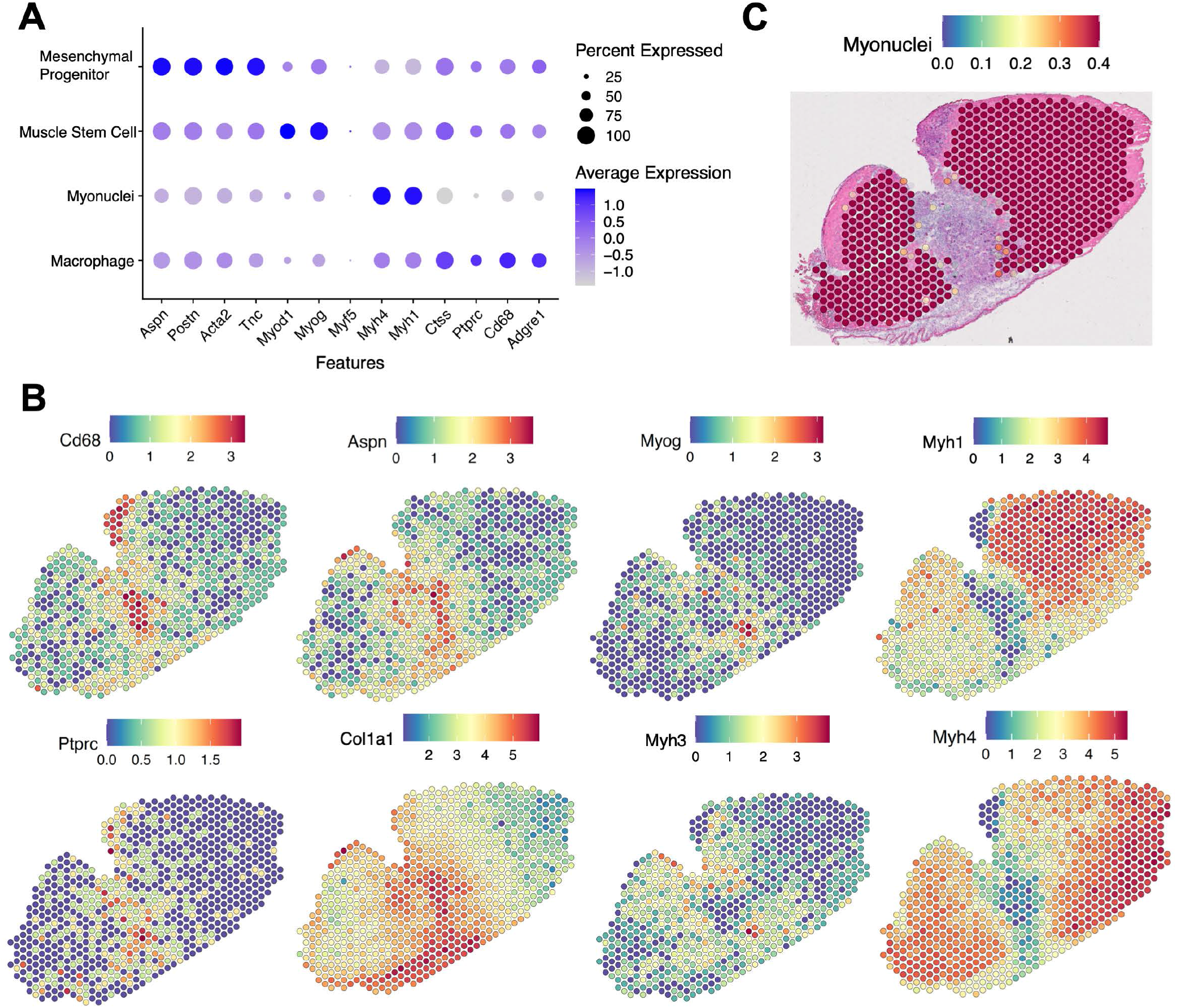
Spatial transcriptomic overlays of genes associated with different cell types after volumetric muscle loss injury. (A) Dotplot showing the expression of known cell-type marker genes in annotated spGEX spots. (B) Overlays of selected cell marker genes are consistent with Seurat label transfer annotations. (C) Spatial overlay of myonuclei prediction scores for each spot.

**Supplemental Figure 3.**
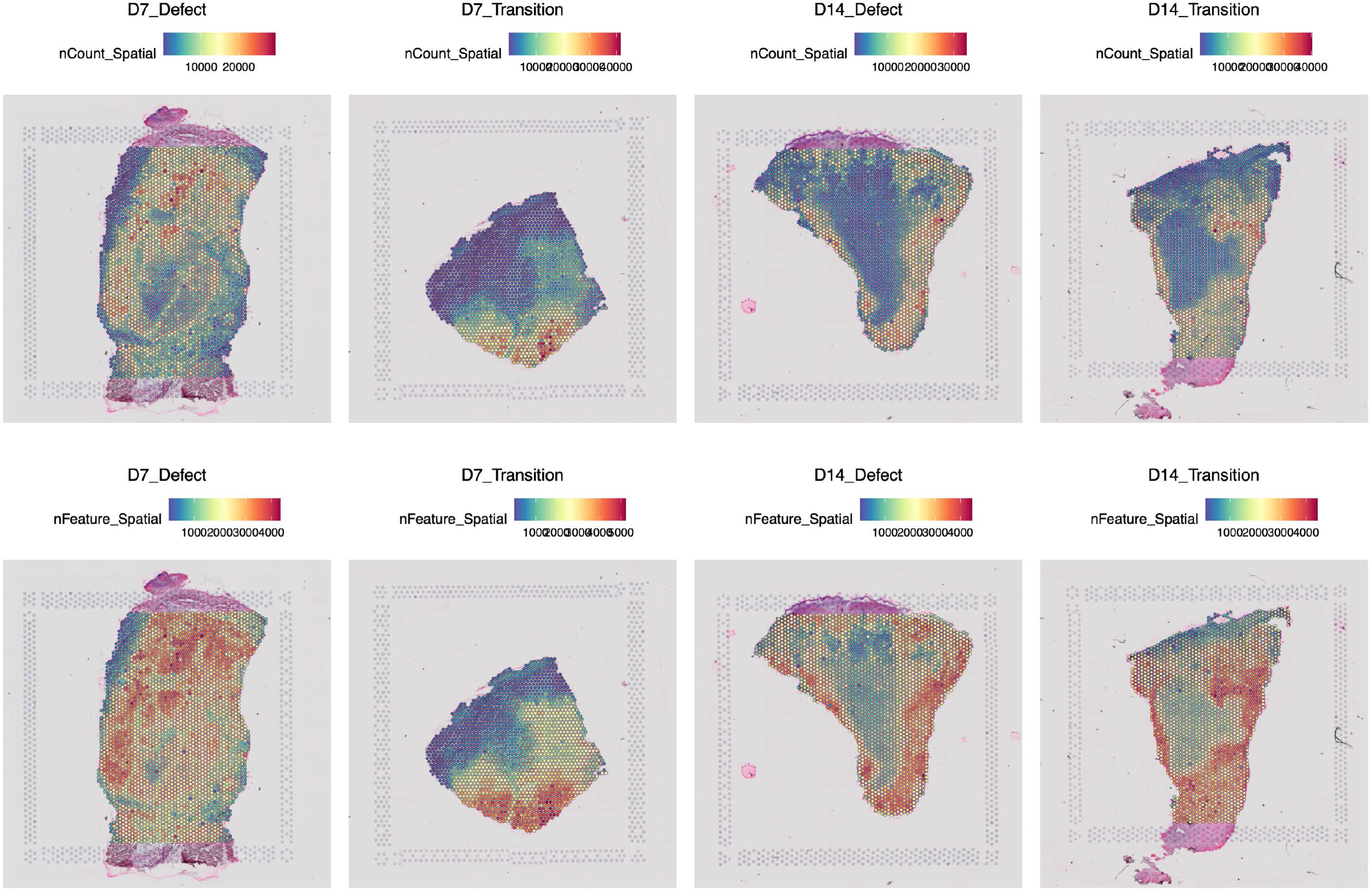
Quality control assessment of spGEX datasets from canines post VML. (A) Overlays of UMIs in each tissue section. (B) Overlays of the number of features identified in each spot.

**Supplemental Figure 4.**
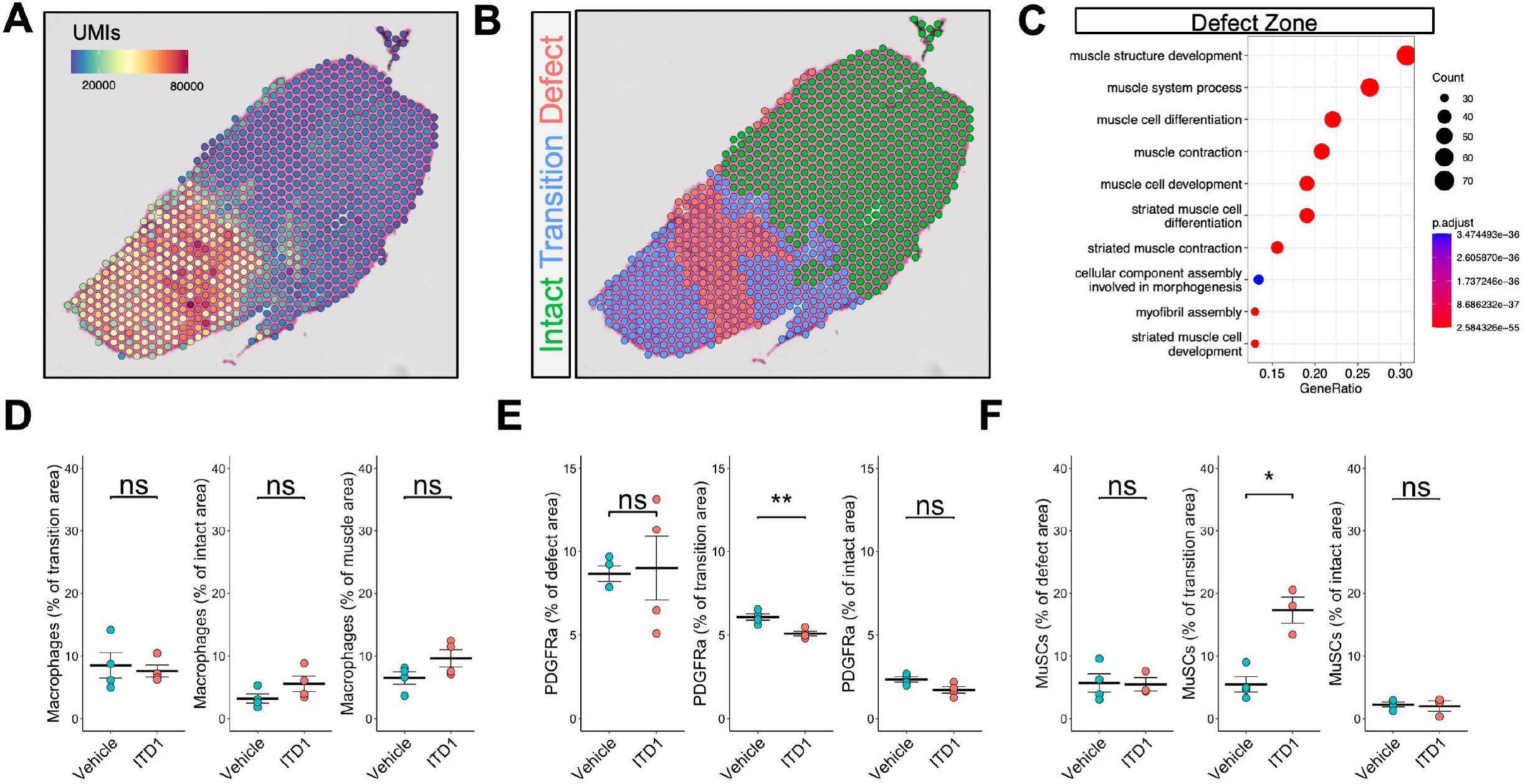
Characterization of ITD1-treated TA VML defects and differences among the transcriptional landscape of the defect zone compared to vehicle controls. (A) Distribution of unique molecular identifiers shows higher read counts at the location of the defect and transitional zones, consistent with untreated tissues. (B) Tissues annotation into the three zones. (C) GOTerm analysis comparing ITD1 vs PBS-treated tissues in the defect zone. The defect zone in ITD1-treated tissues was enriched for terms associated with muscle repair and regeneration, consistent with increased numbers of Pax7+ cells and their progeny in the defects.

## KEY RESOURCES TABLE

| REAGENT or RESOURCE | SOURCE | IDENTIFIER |
| --- | --- | --- |
| **Antibodies** |  |  |
| Rat anti-mouse Ly6G | Abcam | ab25377; RRID:AB_470492 |
| Rat anti-mouse CD68 | BioRad | MCA1957; RRID:AB_322219 |
| Rabbit anti-mouse RFP | Rockland | 600-401-379; RRID:AB_2209751 |
| Chicken anti-mouse GFP | Abcam | ab13970; RRID:AB300798 |
| Rabbit anti-mouse Laminin 1+2 | Abcam | ab7463; RRID:AB_305933 |
| Goat anti-chicken IgY (H+L), Alexa Fluor 488 conjugate | Thermo Fisher | A32931TR; RRID:AB_2866499 |
| Goat anti-rat (H+L), Alexa Fluor 647 conjugate | Thermo Fisher | A21247; RRID:AB_141778 |
| Goat anti-rabbit (H+L), Alexa Fluor 555 conjugate | Thermo Fisher | A21247; RRID:AB_141778 |
| **Chemicals, Peptides, and Recombinant Proteins** |  |  |
| Dispase II (activity $\geq 0.5$ units/mg solid) | Sigma | D4693-1G |
| Collagenase Type II (654 U/mg, non-specific proteolytic activity 487 U/mg) | Life Technologies | 17101015 |
| DMEM, high glucose, pyruvate | Life Technologies | 11995065 |
| Ham's F-10 Nutrient Mix | Life Technologies | 11550043 |
| Fetal Bovine Serum | Life Technologies | 10437028 |
| Normal Goat Serum | Abcam | Ab7481; RRID:AB_2716553 |
| Mouse on Mouse blocking reagent | Vector Labs | MKB-2213 |
| Tissue Plus O.C.T Compound | Fisher Scientific | 23-730-571 |
| Hematoxylin | Ricca Chemical Company | 3530-16 |
| Eosin | EMD-Millipore | 588X-75 |
| Magnesium Sulfate Heptahydrate | Sigma Aldrich | 63138-250G |
| Sodium Bicarbonate | Sigma Aldrich | S5761 |
| SafeClear II | Fisher Scientific | 23-044192 |
| Direct Red 80 | Fisher Scientific | AAB2169306 |
| Picric Acid | Sigma Aldrich | P6744-1GA |
| Glacial Acetic Acid | Sigma Aldrich | BP2401-500 |
| Xylenes | Sigma Aldrich | 534056-4L |
| Permout | Fisher Scientific | SP15-100 |
| Bovine Serum Albumin | Fisher Scientific | BP9703-100 |
| Prolong Diamond | Thermo Fisher | P36965 |
| Tween-20 | Sigma Aldrich | P1379 |
| TritonX-100 | Sigma Aldrich | T8787 |
| Hoechst 33342 | Thermo Fisher | H3570 |
| **Critical Commercial Assays** |  |  |
| QIAshredder | Qiagen | 79656 |
| Qiagen RNeasy Mini Kit | Qiagen | 217084 |
| BioAnalyzer RNA Pico Assay | Agilent | 5067-1514 |
| Tissue Optimization | 10x Genomics | 1000193 |
| Visium Spatial Gene Expression | 10x Genomics | 1000187 |
| **Deposited Data** |  |  |
| spGEX datasets | This Manuscript | GSEXXXXXX |
| **Experimental Models: Organisms/Strains** |  |  |
| C57BL/6J wild-type mice (3 months) | Jackson Labs | Strain 000664;<br>RRID:IMSR_JAX:000664 |
| PDGFRa<sup>EGFP</sup> mice (3 months) | University of Michigan | Strain 007669;<br>RRID:IMSR_JAX:007669 |
| Pax7Cre<sup>ER/+</sup>;Rosa26<sup>dTomato/+</sup> mice (3 months) | University of Michigan | Strain 017763 crossed with strain 007676 |
| **Software and Algorithms** |  |  |
| SpaceRanger v1.3.0 | 10x Genomics | https://support.10xgenomics.com/spatial-gene-expression/software/downloads |
| R v4.1.2 | The R Foundation for Statistical Computing | https://www.r-project.org/<br>RRID:SCR_001905 |
| MATLAB_R2020b | MathWorks | https://www.mathworks.com/products/matlab.html |
| Seurat v4.1.0 | Butler et al. 2019 | https://satijalab.org/seurat/<br>RRID:SCR_007322 |
| CellChat v1.0.0 | Jin 2021 | https://github.com/sqjin/CellChat |
| ggplot2 v3.3.5 | Wickham 2016 | https://ggplot2.tidyverse.org<br>RRID:SCR_014601 |
| dplyr v1.0.8 | Wickham 2016 | https://dplyr.tidyverse.org/<br>RRID:SCR_016708 |
| Tidyverse v1.3.0 |  | https://www.tidyverse.org/<br>RRID:SCR_019186 |
| ImageJ v2.1.0 |  | https://imagej.net/ImageJ<br>RRID:SCR_002285 |
| MAST v1.20.0 | Finak 2015 | https://github.com/RGLab/MAST<br>RRID:SCR_016340 |
| EnhancedVolcano v.1.12.0 | Blighe K. et al. 2019 | https://github.com/kevinblighe/EnhancedVolcano |
| QuPath | Bankhead et al. 2017 | RRID:SCR_018257 |
| **Other** |  |  |
| Bioinformatics analysis code | This manuscript | https://github.com/larouchej/Spatial_VML |

## METHODS

### Animals

C57BL/6 wild-type and PDGFRa^EGFP^ female and male mice were obtained from Jackson Laboratories or a breeding colony at the University of Michigan (UM). Pax7Cre^ER/+^-Rosa26^dTomato/+^ mice were obtained from a breeding colony at UM and administered 5 daily 100uL intraperitoneal injections of 20mg/mL tamoxifen in corn oil. All mice were fed normal chow ad libitum and housed on a 12:12 hour light-dark cycle under UM veterinary staff supervision. All procedures were approved by the University Committee on the Use and Care of Animals at UM and the Institutional Animal Care and Committee and were in accordance with the U.S. National Institute of Health (NIH).

Approval for the procedures conducted on canines in this study was obtained from the University of Pittsburgh Institutional Animal Care and Use Committee. A VML defect was created in the biceps femoris muscle of one female canine (*Canis familiaris)*. Tissue biopsies were collected at multiple time points on days 7 and 14 following injury for spatial transcriptomics analysis.

### Mouse Injury Model

Mice were anesthetized with 1.5% isoflurane and administered 0.1mg/kg buprenorphine in 100μL saline via IP injection. Puralube ointment was applied to both eyes. Hair was removed from the hindlimbs using Nair hair removal cream. The surgical area was sterilized three times with Providone Iodine followed by 70% ethanol. A 0.5cm incision was made in the skin on the anterior side of each tibialis anterior (TA) muscle followed by the removal of a 2mm full thickness muscle section from middle of the TA muscle. The skin was sutured closed using 6-0 proline sutures, which were removed 7 days post-surgery.

### Canine Injury Model

Prior to sedation, the right lateral thigh was shaved. Surgical plane anesthesia was induced with intravenous administration of 2% thiopental sodium, tracheal intubation, and delivery of inhaled isoflurane. The surgical site was prepared by scrubbing with 70% ethyl alcohol, followed by a topical 10% povidone-iodine solution. The greater trochanter and lateral epicondyle of the femur were identified to establish anatomic landmarks. A 10 × 4 cm wide uppercase “I” shape was marked to establish the initial incision point halfway between the greater trochanter and lateral epicondyle. Skin was incised along the outline of the “I” shape, the subcutaneous tissue was bluntly dissected, and the skin was retracted to visualize the biceps femoris. A corresponding 10 cm × 4 cm template was used to delineate the peripheral borders of the planned defect. The muscle tissue and fascia were removed to a depth of 2-3 cm, depending on the anatomy and size of the dog with care taken to avoid cutting through the bottom of the muscle. Hemorrhage was controlled with electrocautery. The overlying skin was removed, with the result that the wound was left exposed to mimic the typical clinical scenario of spontaneous traumatic injury. The upper leg and wound site were covered with sterile dressings. All animals received postoperative antibiotic prophylaxis for the first 5 days (cephalexin, 25 mg/kg) and an analgesic every 8-12 hours for the first 5 days (buprenorphine 0.002 mg/kg). Each animal was observed daily per routine clinical parameters, including temperature, appetite, activity, and ability to bear weight on the operated leg. Animals were fed a high energy, high protein diet (Advanced Protocol High Density Canine Diet; PMI Nutrition LLC, Henderson, CO) and given unlimited access to water.

### Histology

#### Tissue Sectioning

Muscles were simultaneously flash frozen and embedded in optical cutting temperature (OCT) compound according to 10x Genomics Demonstrated Protocol CG000240 Revision D, then stored at −80°C. Serial cross sections were cut using a cryotome at −20°C at the midpoint of the injury and collected on positively charged glass slides (Fisher #12-550-15). Duplicate cross sections from each tissue were placed on each slide.

#### Hematoxylin and Eosin Staining

Slides were submerged in hematoxylin for two minutes followed by two sets of ten quick immersions in distilled water. Slides were then submerged in Scott’s tap water (20.0g magnesium sulfate heptahydrate and 2.0g sodium bicarbonate in 1L tap water) and distilled water for one minute as well as one set of ten immersions in 80% ethanol. After one minute of Eosin immersion slides were immersed ten times in two different 95% ethanol baths and a 100% ethanol bath. Finally, slides were submerged in SafeClear II for one minute, and two drops of Permount were added before placing the coverslip on top. Brightfield images were taken using a motorized Olympus IX83 microscope at 10X magnification and stitched using the Olympus CellSense software to obtain an image of the complete tissue section.

#### Picrosirius Red Staining

Slides were removed from −80°C, thawed to room temperature, and dried for 30 minutes before fixing in 4% paraformaldehyde in PBS for 15 minutes at room temperature. Following fixation, slides were washed 3 times in PBS for 5 minutes each, 2 times in deionized water for 5 minutes each, allowed to dry for 10 minutes at RT, and incubated in 0.5g Direct Red 80 solubilized in 500mL Picric Acid for 1 hour at room temperature. Next, slides were washed twice with acidified water (2.5mL Glacial Acetic Acid in 500mL deionized water) for 5 minutes each followed by 2 washes in deionized water for 5 minutes each. Tissue samples were then dehydrated in a series of ethanol washes (50%, 70%, 70%, 90%, 100%, 100%), incubated in Xylenes twice for 5 minutes each, and mounted with Permount. Brightfield images were taken using a motorized Olympus IX83 microscope at 10X magnification and stitched using the Olympus CellSense software to obtain an image of the complete tissue section. Images were converted to RGB stack and converted to greyscale. To calculate collagen fraction, the green channel was automatically thresholded and measured using ImageJ, then divided by the surface area of the tissue section based on thresholding the red channel to include only the tissue. Two or three sections per tissue were stained and quantified, then averaged to get the percentage of collagen for each tissue.

#### Immunofluorescent Staining

Immunofluorescence staining was performed as previously reported(1). For RFP and immune stains, slides were removed from −80°C, thawed to room temperature for 30 minutes, then fixed in ice-cold 100% acetone at −20°C for 10 minutes. Following fixation, slides were air-dried for 10 minutes at RT, tissue sections were circled with Hydrophobic Barrier PAP Pen and allowed to dry at room temperature. Tissue sections were the rehydrated in PBS for 5 minutes at RT, then blocked for 1 hour at room temperature using 10% normal goat serum (NGS) in PBS. After blocking, slides were incubated overnight at 4°C in a solution containing primary antibodies (rabbit anti-laminin 1+2 (1:500 dilution), rat anti-laminin 2 alpha (1:1000), rat anti-CD68 (1:50), rabbit anti-RFP (1:100)) diluted in 10% NGS. Primary antibodies were then washed three times for 5 minutes with PBS at room temperature. Secondaries (1:500 dilution) and Hoescht 33342 (1:500 dilution) were added in PBS and incubated for 1 hour at RT in the dark. After incubation, slides were washed three times with PBS and a coverslip was mounted using Prolong Diamond florescent mounting medium. For GFP stain, slides were warmed to room temperature, tissue sections circled in PAP Pen, re-hydrated in PBS three times for 5 minutes, blocked for 30 minutes at RT in MOM blocking reagent, and incubated overnight at 4°C in primary antibodies (1:1000 GFP and 1:500 anti-laminin 1+2) diluted in MOM Protein Concentrate. Following primary incubation, slides were washed 3 times for 5 minutes with PBS and incubated in secondary antibodies (AF488 anti-chicken 1:250, AF555 anti-Rabbit 1:500, 1.5uL/mL Hoescht 33342) diluted in MOM protein concentrate for 1 hour at RT. Finally, slides were washed three times for 5 minutes in PBS, then mounted with Prolong Diamond. Slides were allowed to dry overnight, then stored at 4°C until imaging. Images were acquired with a Nikon A1 confocal microscope equipped with Colibri 7 solid state light source and pseudo colored using ImageJ, then quantified using MATLAB.

#### Automated Image Analysis with QuPath

Samples for histologic analysis were obtained from independent animals not sampled for spatial transcriptomics. Following euthanasia, the full thickness of the entire VML defect was harvested en bloc and fixed in 10% neutral buffered formalin. Post fixation, biopsies from the VML defect and transition zone between the wound bed and wound margin were obtained using 6-8 mm diameter biopsy punches. The specimens were trimmed, embedded in paraffin, and sectioned at 5 μm prior to staining with Masson’s Trichrome. Slides were imaged using a Motic Easy Slide Scanner at a 40X magnification for automated analysis using QuPath(2). QuPath, an open-source image analysis software, was then used to classify regions within the scanned images. QuPath was used to identify healthy muscle, necrotic muscle, granulation tissue, and connective tissue by training the artificial neural network with a library of samples from canine VML defects ranging from 2-42 days post injury. Manual segmentation of scab/debris at the superficial edge of the tissue was performed, and QuPath was then used to classify the remaining tissues within the image. A colored mask was then created to enable visualization of the distinct tissue regions within the sample. Analysis was limited to the most superficial 6 mm of the wound bed, representative of the active wound zone, and to be representative of the tissue biopsies which were collected for spatial transcriptomic analysis.

### Spatial Gene Expression Sequencing

#### Sample Preparation and Sequencing

Tissues (mouse and canine) were simultaneously flash-frozen and OCT embedded as described above. RNA quality for each tissue was assessed using the QIAshredder and RNeasy Mini kit for RNA extraction according to manufacturer’s protocols followed by BioAnalyzer RNA Pico assay and tissues with RNA integrity values above 7 were used for spatial gene expression (spGEX) profiling. Permeabilization times were determined for each timepoint on mouse and canine VML-injured tissues using the 10x Genomics Tissue Optimization kit and corresponding protocol. Canine tissues were permeabilized for 24 minutes, and mouse tissues were permeabilized for 18 minutes. SpGEX profiling was performed using the 10x Visium platform according to manufacturer’s instructions and sequenced on a NovaSeq 6000.

#### Data Processing and Analysis

Manual image alignment was performed using 10x Loupe Browser. 10x SpaceRanger v1.3.0 software’s mkfastq and count command were run with default parameters and aligned to the mm10-2020-A or the CanFam3.1 v97 genome for mouse and canine datasets, respectively. Tissues were annotated into zones using Loupe Browser v6. HDF5 matrix files and zone annotations were imported into R (https://www.r-project.org/) using the Seurat v4 package(3). Count data were normalized using SCTransform, followed by PCA dimension reduction and clustering using default parameters and the first 30 dimensions. GOTerm analysis was performed on scaled, normalized data using FindAllMarkers followed by ClusterProfiler’s enrichGO function on upregulated genes. Mitochondrial and ribosomal genes were removed prior to GOTerm enrichment analysis. For mouse datasets, spot annotation with cell-types was performed using Seurat label transfer with a matched single cell RNA-Seq reference generated from 3mm quadricep defects at 7 days post injury(4). Ligand-receptor analysis was performed on the single cell RNA-Seq dataset used for label transfer using CellChat(5). Gene modules were generated using Seurat’s AddModuleScore function for overlays. Differential gene expression comparing the defect and transition zones between treated and untreated tissues was performed using MAST(6). Plots were generated using ggplot2, EnhancedVolcano, and Seurat.

### Statistics

Experiments were repeated at least twice. Bar graphs show mean ± standard error. Statistical analysis was performed in R using two-sample Student’s t-test assuming normal distribution and equal variances or one-way ANOVA, as specified in figure captions. All statistical tests performed were two-sided. Outliers were determined using the Tukey’s fences method with k = 1.5 and removed from further analysis. P-values less than 0.05 were considered statistically significant.

